# A germline-specific reporter reveals conserved primordial germ cell developmental dynamics between jawless and jawed fish species

**DOI:** 10.64898/2026.09.23.753524

**Authors:** Sreeja Sarasamma, Yu-Wen Chung Davidson, Tyler J. Buchinger, Jiahao Song, Weiming Li

## Abstract

Primordial germ cells (PGCs) are the stem cell lineage responsible for transmitting genetic information across generations and are therefore central to studies of germline development, reproduction, and evolution. However, a lack of molecular tools in jawless vertebrates such as lampreys and hagfish has limited comparative insights into the evolutionary origins of vertebrate germline cells. Here, we report the development of the first germline-specific reporter (GSR) in the sea lamprey (*Petromyzon marinus*), Tg(piwil1: egfp-UTRnanos1), a *piwil*1-based germline reporter that is driven by the sea lamprey piwil1 regulatory region that directs Green Fluorescent Protein (GFP) expression in early PGCs. Promoter activity was validated by robust GFP expression in PGCs of zebrafish embryos co-injected with Tol2 transposase mRNA. Injection of the construct into sea lamprey embryos resulted in early, persistent, and gonad-localized GFP-positive cells. These results demonstrate stable germline-specific transgene expression in a jawless vertebrate and suggest deep evolutionary conservation of germline regulatory mechanisms between jawless and jawed vertebrates. Furthermore, the piwil1-based germline reporter line enabled lifetime visualization of zebrafish germline cells, revealing that increased PGC abundance during embryogenesis biases toward female sexual differentiation. Together, this study establishes the first GSR system in a jawless vertebrate and provides a useful platform for investigating germline biology and vertebrate germline evolution.

## Introduction

Primordial germ cells (PGCs) constitute the lineage that transmits genetic and epigenetic information across generations, placing them at the core of vertebrate development and evolution (1–3) (Yu et al.,2025; Heard, E. and Martienssen, R. A. 2014; Weissman, A. 1892). As the earliest committed germline precursors, PGCs are essential for reproduction, fertility, and inheritance. Studies in classical model organisms, including mice, zebrafish, and Xenopus, have uncovered conserved molecular mechanisms governing PGC specification, migration, and differentiation (4–8)(Whittle CA, Extavour CG.,2017; Barton LJ et al.,2024; Mikedis MM, Downs KM,2014; Bertho S et al.,2021; Johnson & Alberio, 2015). However, this understanding remains largely restricted to jawed vertebrates, leaving key questions unresolved regarding how germline regulatory programs evolved during early vertebrate diversification. Recent advances, including the derivation of human PGC-like cells and high-resolution profiling of migratory germ cells, have highlighted the dynamic regulatory landscape of germline development (9, 10)(Esfahani et al., 2024; Jaszczak et al., 2025). Despite this progress, our understanding of germline regulatory programs remains largely focused on jawed vertebrates, while functional tools to directly visualize and interrogate germline lineages in jawless vertebrates remain limited. This gap constrains our ability to determine how germline regulatory programs evolved during early vertebrate diversification and to reconstruct ancestral mechanisms of germline development.

Cyclostomes (lampreys and hagfishes) provide a critical phylogenetic perspective for understanding the evolution of germline specification (14) (Marlétaz F, et al.,2024). Lampreys occupy a pivotal phylogenetic position as one of the earliest diverging vertebrate lineages, providing a unique opportunity to investigate the evolutionary origins of vertebrate developmental mechanisms. Comparative studies between cyclostomes and jawed vertebrates have revealed both conserved and lineage-specific features of genome organization, development, and reproductive biology. Understanding germline specification and regulation in lampreys, therefore, provides important insights into how vertebrate germline systems evolved during early vertebrate diversification (15)(Hughes LC et al.,2025).

Two major strategies underlie PGC specification in vertebrates. In the inductive mode, as observed in mammals, extracellular signaling pathways such as BMP and WNT direct pluripotent embryonic cells toward germline fate through transcriptional regulators, including PRDM1/BLIMP1, PRDM14, and TFAP2C (11)(Makela JA.,2019). By contrast, in preformation, common among teleosts and amphibians, maternal germ plasm enriched in determinants such as vasa and nanos is asymmetrically inherited to establish PGC identity (12) (Nishimura T, Fujimoto T, 2025). Comparative analyses suggest that inductive specification may represent the ancestral vertebrate state, whereas preformation likely evolved secondarily; however, functional data from jawless vertebrates remain insufficient to test this hypothesis (13)(Hansen & Pelegri, 2021). Investigations of germline specification in lampreys have been limited to descriptive histology or in situ detection of germline transcripts. Currently, there are no stable germline-specific transgenic tools that enable direct identification, visualization, and lineage analysis of PGCs in jawless vertebrates such as the sea lamprey. This limitation has restricted experimental access to the PGC lineage in these organisms. Recent advances in genome engineering technologies, particularly CRISPR-based platforms, have expanded the potential for functional genomics in non-model vertebrates, including the sea lamprey (16)(Sarasamma & Li, 2026). However, tools enabling direct visualization and lineage analysis of germline cells in jawless vertebrates remain largely absent. Technical barriers are further compounded by programmed genome rearrangement (PGR) in sea lamprey, where ∼20% of the genome is eliminated from somatic lineages early in embryogenesis (17, 18)(Smith et al., 2009; Timoshevskiy et al., 2016).

Recent single-cell transcriptomic and multi-omics studies have begun to resolve the molecular heterogeneity and regulatory landscapes of migratory PGCs, but comparable datasets from cyclostomes remain scarce. This limitation leaves a significant gap in our understanding of germline regulatory networks at the base of the vertebrate lineage(10) (Jaszczak RG et al.,2025). Recent genomic analyses have further revealed that the sea lamprey germline genome contains specialized germline-restricted chromosomes and unique structural features associated with PGR, highlighting the distinctive evolutionary architecture of the cyclostome germline (19)(Timoshevskaya et al., 2023). Earlier studies demonstrated that large portions of the genome are selectively eliminated from somatic cells during development, while germline-specific sequences are retained to support reproductive function (20)(Smith et al., 2012). Recent analyses reveal that many germline-restricted genes are testis-biased and may influence reproductive biology (21) (Yasmin et al., 2022). This unusual genome architecture complicates reporter design but also presents opportunities to exploit germline-specific regulatory elements (19) (Timoshevskaya et al., 2023).

Beyond its evolutionary significance, germline biology is closely linked to sex determination and sexual differentiation. Early observations in cyclostomes suggested that variation in PGC number may influence reproductive capacity, as PGC abundance was correlated with fecundity in developing lamprey gonads (22)(Hardisty & Cosh, 1966). In zebrafish, germ cell abundance during early development strongly biases sex outcome: individuals with higher PGC numbers tend to differentiate as females, whereas depletion promotes male development or sterility (23–25) (Tzung KW et al.,2015; Dai et al., 2015; Fontana et al., 2023). Germline proliferation and meiotic entry have been shown to diverge in presumptive sexes (26) (Pan et al., 2022), and epigenetic reprogramming of the germline methylome is implicated in sex transitions (27)(Wang et al., 2021). These findings suggest that PGCs actively contribute to sexual differentiation rather than merely responding to somatic developmental cues, raising the question of whether such interactions reflect an ancient vertebrate feature or a gnathostome-specific innovation remains unresolved.

Across vertebrates, germline development is regulated by conserved molecular pathways that maintain germline identity and genome integrity. The PIWI-piRNA pathway and germline-associated genes such as *piwil1*, *nanos*, and *vasa* play important roles in germline maintenance and have been widely used to identify and track primordial germ cells (PGCs) in both jawless and jawed vertebrates (28)(Ku HY, Lin H., 2014)(29)(Zawalska M, Tarnowski M.,2025).(30)(Juliano C et al.,2011).(31, 32)(Thomson & Lin, 2009;Iwasaki et al., 2015)(33)(Houwing S, et al.,2007).(34) (Suzuki H et al.,2010)(35, 36) (Mishima Y et al.,2006;Jin YN et al.,2018). The conserved germline expression of *piwil1* provides a strong basis for transcriptional targeting, while the *nanos* 3′ untranslated region (3′ UTR) can enhance germline specificity through post-transcriptional regulation (37)(Lee CY, et al.,2017)(13)(Hansen & Pelegri, 2021).(36, 38)(Jin YH et al.,2019; Chu WK et al.,2025)(39) (Kaufman OH, Marlow FL.,2016).

In vertebrates, PIWI proteins are highly enriched in germ cells and have been widely used as markers of germline identity. The conserved germline expression pattern of *Piwil1*, therefore, provides a promising regulatory framework for developing GSR systems (33)(Houwing S, et al.,2007). Despite the conserved roles of PIWI proteins in germline regulation, the functional activity of sea lamprey piwil1 regulatory elements has not been experimentally characterized. In addition to their role in maintaining germline identity, germline regulatory genes such as *piwil1*, *nanos*, and *vasa* have been widely used as molecular markers to visualize PGCs in diverse vertebrate models (36, 38)(Jin YH et al.,2019; Chu WK et al.,2025). Transgenic reporter systems based on germline regulatory elements have enabled direct observation of PGC specification, migration, and colonization of the developing gonads in teleost fishes and other vertebrates. These tools have greatly advanced our understanding of germline lineage dynamics and reproductive biology by allowing *in vivo* tracking of germ cell populations throughout development (39) (Kaufman OH, Marlow FL.,2016). Because cyclostomes diverged from the gnathostome lineage more than 500 million years ago, lampreys provide a valuable system for investigating conserved and potentially ancestral features of vertebrate germline biology. However, the limited availability of germline-specific molecular tools in lampreys has constrained experimental investigation of these processes.

We hypothesized that regulatory elements of the sea lamprey *piwil1* gene could be used to establish a germline-specific reporter that faithfully marks PGCs and reveals conserved features of germline development across jawless and jawed vertebrates. To test this hypothesis, we combined the sea lamprey *piwil1* regulatory region with the *nanos1* 3′ UTR to develop the first germline-specific reporter (GSR) system in the sea lamprey, Tg(piwil1-UTRnanos1). We characterized reporter expression and PGC dynamics across developmental stages in sea lamprey and zebrafish and performed a cross-species functional analysis of the sea lamprey *piwil1* regulatory region in zebrafish. We further investigated the relationship between PGC abundance and sex differentiation using lifetime germline labeling. Collectively, this work establishes a molecular toolkit for visualizing and quantifying germline cells in sea lamprey and provides experimental evidence for conserved germline regulatory activity across jawless and jawed vertebrates. The reporter system provides a foundation for investigating germline development, PGC dynamics, and their relationship to sexual differentiation in sea lamprey, with potential applications to future studies of germline biology and genetic approaches for managing invasive sea lamprey populations in the Great Lakes.

## Materials and Methods

### Experimental Animals and Maintenance

Wild-type zebrafish (*Danio rerio*) and sea lamprey (*Petromyzon marinus*) were used in this study. All experimental procedures were conducted in accordance with institutional guidelines and were approved by the Institutional Animal Care and Use Committee (IACUC) at Michigan State University (PROTO202200326, PROTO202200296, PROTO202500268, PROTO202500267). Adult zebrafish were obtained from the Zebrafish International Resource Center (ZIRC, Eugene, USA) and maintained in a recirculating aquatic system under standard laboratory conditions at 28.5 °C with a 14 h light / 10 h dark photoperiod. Fish were fed twice daily with a combination of commercial dry feed and live *Artemia nauplii*. Zebrafish husbandry and breeding procedures were carried out following established laboratory protocols (40) (Westerfield, 2000). For embryo collection, breeding pairs were placed in spawning tanks the evening before spawning, and fertilized embryos were collected shortly after fertilization. Embryos were rinsed and maintained in E3 embryo medium at 28.5 °C in Petri dishes and staged according to established zebrafish developmental criteria. Embryos were monitored regularly under a Zeiss Stemi 508 stereomicroscope (Carl Zeiss Microscopy LLC, Thornwood, NY), and unfertilized or abnormal embryos were removed to ensure healthy development. Adult sea lampreys (*Petromyzon marinus*) were obtained from the US Geological Survey Hammond Bay Biological Station (Millersburg, MI, USA). Sea lamprey embryos were generated from spawning adults collected from traps operated by the US Fish and Wildlife Service and Fisheries and Oceans Canada in tributaries of the Great Lakes region during the natural spawning season. Fertilized eggs were produced through controlled crosses and maintained in aerated freshwater under laboratory conditions at approximately 16-18°C, following established lamprey husbandry and embryological procedures (41, 42)(Piavis, 1961; McCauley et al., 2015). Embryos were incubated in fresh water (Aquaneering, San Marcos, CA) and monitored daily, and dead or unfertilized embryos were removed to maintain water quality. Developing embryos were examined using a Zeiss Stemi 508 stereomicroscope, and embryos at the appropriate developmental stages were selected for microinjection and fluorescence imaging experiments.

### Generation of a piwil1-Based Germline Reporter Construct

We generated a germline-specific reporter construct using the HLC vector, a Tol2-based plasmid backbone containing Tol2 long terminal repeats (LTRs) and I-SceI meganuclease recognition sites to enable genomic integration. The original HLC vector, which contained a *c-fos* minimal promoter driving enhanced green fluorescent protein (EGFP) followed by an SV40 polyadenylation signal, was provided by the Stowers Institute for Medical Research. The *c-fos* promoter was excised using EcoRI and NcoI restriction enzymes, and the resulting backbone, retaining the EGFP coding sequence, was purified and used for subsequent cloning. The *c-fos* regulatory element was replaced with the approximately 3.8-kb sea lamprey *piwil1* upstream regulatory region, followed by the *nanos1* 3′UTR, to generate the Tg(piwil1:EGFP–UTRnanos1) germline-specific reporter construct.

A ∼3.8 kb upstream regulatory region of the sea lamprey *piwil1* gene was amplified from genomic DNA and cloned upstream of the EGFP coding sequence to drive germline-specific transcription, consistent with the conserved role of PIWI proteins in germline maintenance (31)(Thomson & Lin, 2009). To further restrict expression to germ cells, the 3′ untranslated region (3′UTR) of *nanos1* was inserted downstream of EGFP using compatible restriction sites (SacI or KpnI), replacing or supplementing the SV40 polyadenylation signal. The *nanos1* 3′UTR confers post-transcriptional stabilization of transcripts in primordial germ cells, thereby restricting reporter expression to the germline lineage (43) (Köprunner et al., 2001). The final expression cassette (*piwil1*:EGFP–UTRnanos1) was flanked by Tol2 elements, generating a germline-specific reporter construct. All constructs were assembled using standard restriction-ligation cloning and validated by diagnostic restriction digestion and Sanger sequencing before embryo microinjection.

### Construct Validation

Plasmid constructs were verified by diagnostic restriction digestion and Sanger sequencing before embryo microinjection. Following injection, reporter expression was assessed by fluorescence microscopy, and embryos exhibiting germline-associated GFP fluorescence were selected for subsequent analysis.

### Microinjection of Sea Lamprey and Zebrafish Embryos

Fertilized sea lamprey embryos were obtained using standard *in vitro* fertilization procedures and maintained in aerated, filtered freshwater at approximately 16-18 °C until microinjection. For zebrafish, fertilized embryos were collected shortly after natural spawning and maintained under standard conditions. Reporter plasmid DNA was purified using an endotoxin-free plasmid preparation kit (QIAprep Spin Miniprep Kit; Qiagen) and diluted to a final concentration of 20-40 ng µL⁻¹ in injection buffer. Approximately 1-2 nL of plasmid solution was microinjected into one-cell stage embryos using a pneumatic microinjector (PV850 Pneumatic PicoPump, World Precision Instruments, Sarasota, FL, USA) under a Zeiss Stemi 508 stereomicroscope. To facilitate genomic integration, Tol2 transposase mRNA or I-SceI meganuclease was co-injected with the plasmid DNA, depending on the integration strategy. Following injection, embryos were incubated under standard conditions (16-18 °C for sea lamprey; 28.5 °C for zebrafish) and monitored throughout embryonic development.

### Survival Analysis

Embryo survival following microinjection was monitored to determine whether introduction of the reporter construct affected developmental viability. For both zebrafish and sea lamprey,100 fertilized embryos were established as the initial cohort for each injected and uninjected control group and maintained under identical conditions and monitored throughout early development. Embryos were monitored longitudinally at defined developmental stages, and survival was recorded relative to the initial cohort size. Survival was recorded at defined developmental stages and expressed as the percentage of surviving embryos relative to the initial number of fertilized embryos. Survival curves were generated using the Kaplan-Meier method, and differences between groups were assessed using the log-rank (Mantel-Cox) test. Statistical analyses were performed using GraphPad Prism, and significance was defined as *p* < 0.05 (44) (Kaplan EL, Meier P).

### Fluorescence Imaging and Confocal Microscopy

Injected embryos were screened for EGFP fluorescence at multiple developmental stages using an inverted fluorescence microscope (ECHO Revolution; BICO Company, San Diego, CA, USA). Cells exhibiting localized GFP fluorescence consistent with primordial germ cell (PGC) distribution were identified based on their characteristic anatomical location and migratory behavior during embryogenesis, as previously described (45) (Raz, 2003). For high-resolution imaging, embryos and larvae were anesthetized with 65 mg/L tricaine (MS-222, Sigma-Aldrich, St. Louis, MO, USA), mounted in 1% ultra-pure low-melting-point agarose, and imaged using confocal and multiphoton (two-photon) microscopy (Leica Microsystems, Inc., Buffalo Grove, IL, USA). Z-stack images were acquired and processed using standard image analysis software to visualize the three-dimensional distribution of labeled germ cells. Transgenic larvae at defined developmental stages were dissected to expose the gonadal region. Samples were fixed in 4% methanol-free paraformaldehyde at 4 °C overnight, washed with 0.1M phosphate buffer saline (PBS) containing 0.1% Triton X-100, and stained with phalloidin Alexa Fluor 568 to visualize F-actin and DAPI for nuclear labeling. Samples were equilibrated in 50% glycerol-PBS before imaging. High-resolution imaging was performed using a laser-scanning confocal microscope (Leica Stellaris 8 DIVE, Leica Microsystems) equipped with a 25×/1.1 NA water immersion objective. Z-stack images were acquired at 0.5 µm intervals and processed using Fiji/ImageJ to reconstruct three-dimensional tissue organization (46) (Schindelin et al., 2012).

### Immunofluorescence and Immunohistochemistry (IF&IHC) for Germ Cell Validation

Germline identity was validated using IF & IHC with established germ cell markers. Sequential double-staining was performed to confirm antibody specificity while minimizing antigen damage from repeated retrieval steps. Primary antibodies included rabbit anti-DDX4/VASA (1:1000; AB4330, Sigma-Aldrich,St. Louis, MO, USA) and rabbit anti-NANOS1 (1:1000; PRS4685, Sigma-Aldrich, St. Louis, MO, USA). Sea lamprey larvae were collected at multiple developmental stages (5-15 dpf, n = 20 per stage; 20 and 30 dpf, n = 120 per stage). Tissue sections were deparaffinized in xylene (2 × 15 min), rehydrated through a graded ethanol series (100% to 50%), and washed in TBST buffer (50 mM Tris-HCl, 150 mM NaCl, pH 7.4, 0.1% Tween-20). Antigen retrieval was performed using antigen retrieval buffer (Proteintech PR30001, 1:50 dilution) at 100 °C for 10 min, followed by quenching of endogenous peroxidase activity with 3% hydrogen peroxide. Samples were stained with phalloidin Alexa Fluor 568 to visualize F-actin and counterstained with DAPI (1 µg/mL) for nuclear labeling. Following staining, samples were equilibrated in 50% glycerol-PBS before imaging. (47)(Hsu et al., 1981).

### Germ Cell Imaging and Quantification

Transgenic zebrafish embryos expressing Tg(piwil1: EGFP-UTRnanos1) were analyzed at key developmental stages 24 hpf, 48 hpf, 5 dpf, 10-15 dpf, and 60-90 dpf. Embryos and larvae were anesthetized with tricaine, mounted in low-melting-point agarose, and imaged by confocal microscopy. GFP-positive primordial germ cells (PGCs) were identified based on characteristic localization and migration. Early stages (24-48 hpf) were used to assess germ cell migration and gonadal colonization. At 5 dpf, PGC localization and abundance were evaluated; PGCs were quantified in individual embryos, and embryos were classified into PGC-low and PGC-high groups based on the median PGC count of the analyzed population; embryos with counts below the median were classified as PGC-low, whereas those with counts at or above the median were classified as PGC-high. Germ cell persistence within the gonadal region was examined at 10-15 dpf. Fish were raised to adulthood, and sex was determined at 60-90 dpf based on gonadal morphology. Sex ratios were calculated as the proportion of males and females in each group (48).

### Statistical Analysis

Statistical analyses were performed to compare germ cell counts and sex ratios between experimental groups. Differences in proportions were assessed using appropriate statistical tests, including chi-square or Fisher’s exact test. Results were considered statistically significant at p < 0.05. Data visualization and statistical analyses were performed using GraphPad Prism.

## Results

### Evolutionary Conservation of PIWIL1 Across Vertebrates

The sea lamprey PIWIL1 sequence clustered within the vertebrate PIWI family and was closely related to PIWIL1 sequences from teleosts, including killifish and zebrafish (Fig. 1A). To examine the evolutionary relationship of sea lamprey *piwil1* to homologous vertebrate sequences, we compared PIWIL1 protein sequences from representative vertebrates, including human, mouse, chicken, killifish, zebrafish, and sea lamprey. Multiple sequence alignment further identified conserved amino acid residues within functional PIWI domains across the analyzed vertebrate sequences (Fig. 1B). These conserved sequence features support the evolutionary conservation of PIWIL1 protein architecture and provided a rationale for investigating the activity of the sea lamprey *piwil1* upstream regulatory region in a germline reporter system.

**Figure 1.**
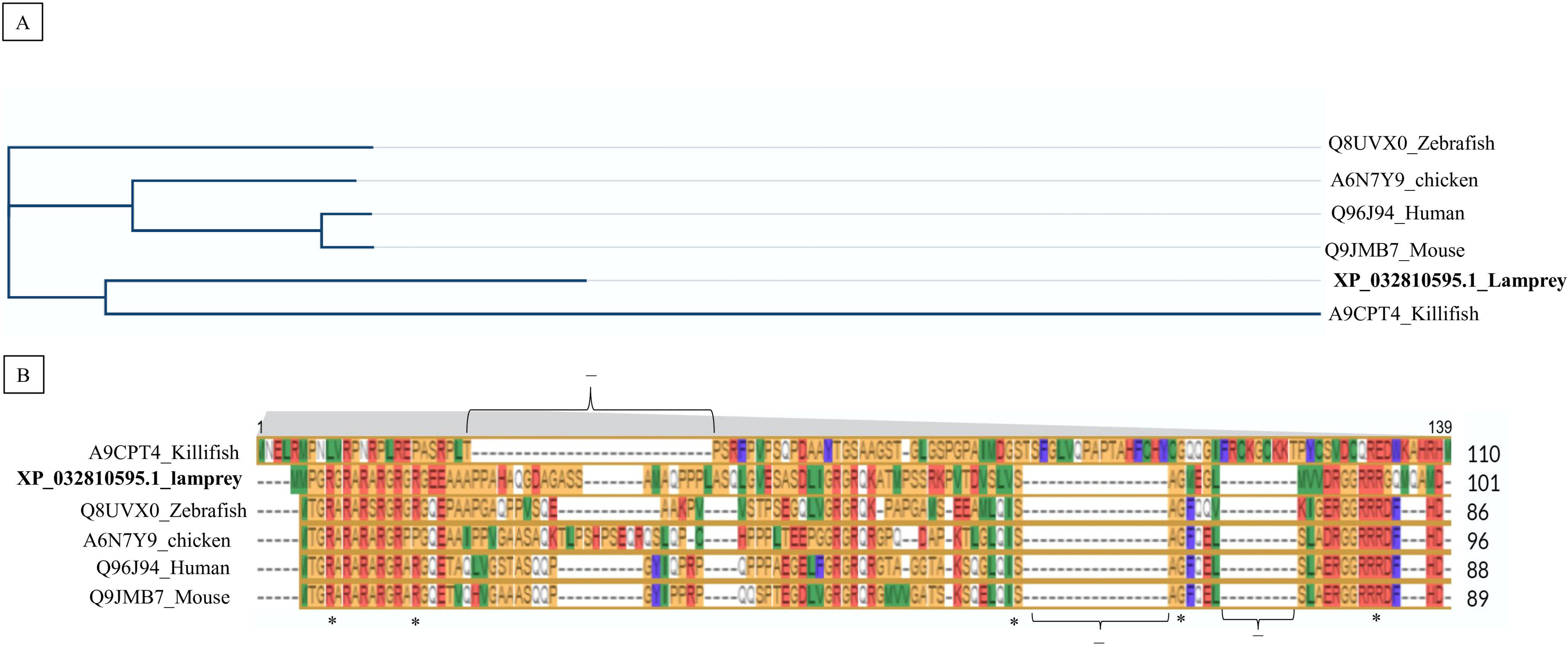
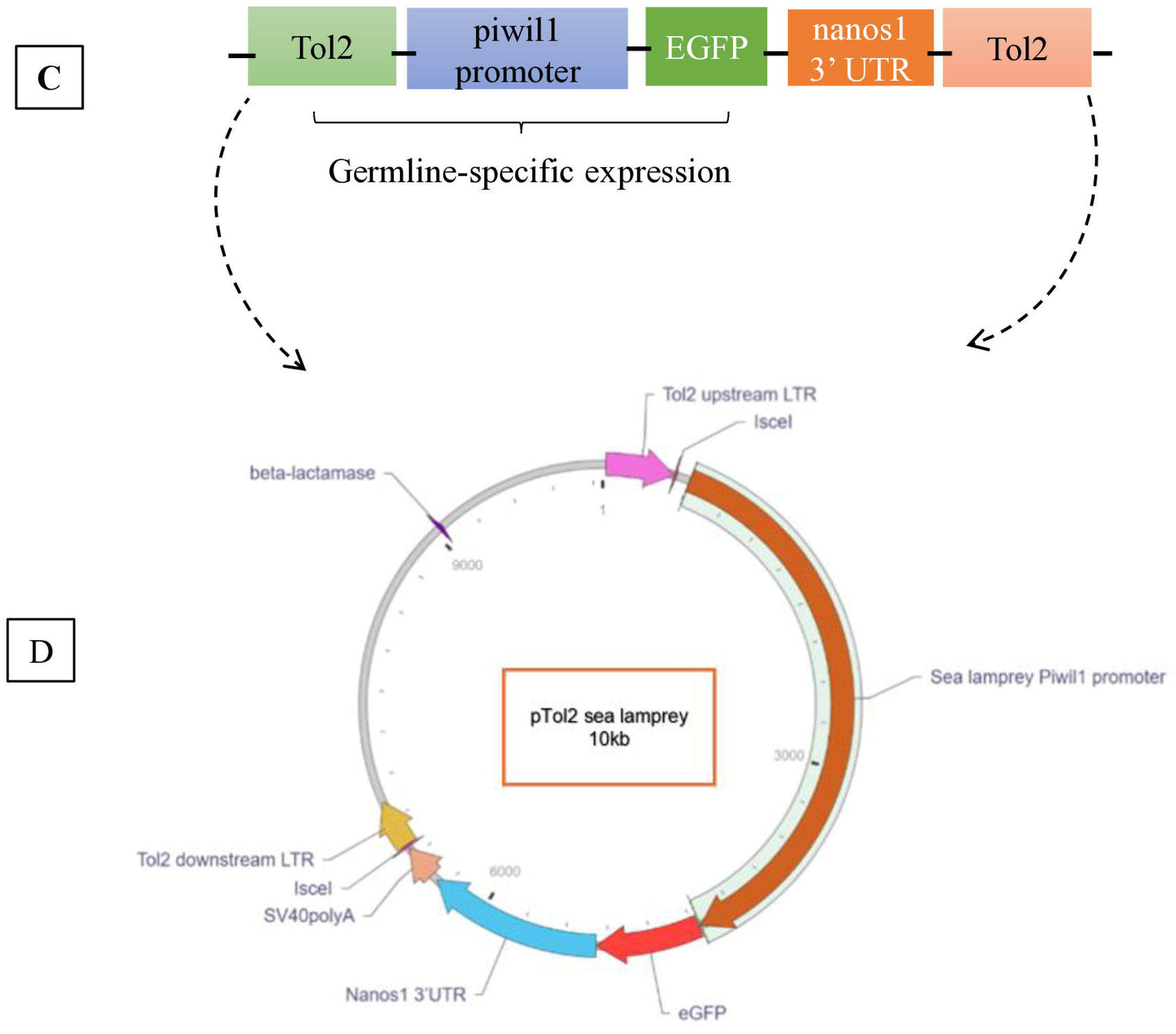
Evolutionary conservation of PIWIL1 across vertebrates. (A) Phylogenetic analysis of PIWIL1 protein sequences from representative vertebrates, including human, mouse, chicken, zebrafish, and sea lamprey. The tree shows that the sea lamprey PIWIL1 clusters within the vertebrate PIWI clade, supporting deep evolutionary conservation of this protein family. (B) Multiple sequence alignment of PIWIL1 proteins highlighting conserved amino acid residues across species. Conserved regions corresponding to functional PIWI domains are evident, indicating preservation of key molecular features associated with germline regulation. Identical and highly conserved residues are highlighted, with gaps introduced to optimize alignment. Accession numbers for sequences used in the analysis are provided alongside species names. (C) Schematic representation of the Tg(piwil1: EGFP-UTRnanos1) germline reporter construct. The reporter consists of the sea lamprey piwil1 promoter driving EGFP expression, followed by the nanos1 3′ untranslated region (3′UTR) to confer post-transcriptional germline specificity. The expression cassette is flanked by Tol2 transposable elements to facilitate stable genomic integration. (D) Functional strategy of the germline reporter. The piwil1 promoter directs transcription in germline cells, while the nanos1 3′UTR restricts and stabilizes reporter expression within primordial germ cells (PGCs), resulting in germline-specific EGFP fluorescence for visualization and lineage tracing during vertebrate development. * Identical amino acid residues across all sequences. - Alignment gaps introduced to maximize sequence similarity.

### Design and Assembly of a Germline-Specific Reporter Construct

The Tg(piwil1:EGFP–UTRnanos1) reporter construct was successfully assembled and sequence-validated for visualization of germline cells (Fig. 1C, D). The construct contained an approximately 3.8-kb sea lamprey *piwil1* upstream regulatory region positioned upstream of the EGFP coding sequence, followed by the *nanos1* 3′UTR, with the expression cassette flanked by Tol2 transposon elements for genomic integration (Fig. 1C, D). Restriction enzyme analysis produced the expected fragment pattern, consistent with the predicted organization of the reporter cassette. Sanger sequencing further confirmed the integrity and expected sequence of the cloned *piwil1*-EGFP-*nanos1* reporter region. The sequence-verified construct was subsequently used for embryo microinjection and analysis of GFP-positive germ cells.

### Survival of Injected Embryos

The Tg(piwil1-UTRnanos1) reporter construct did not significantly impair embryonic survival in either zebrafish or sea lamprey (Fig. 2). In zebrafish, injected embryos exhibited survival rates comparable to uninjected controls throughout early development (Fig. 2A). Similarly, sea lamprey embryos injected with the reporter construct showed normal developmental progression and survival rates comparable to uninjected embryos (Fig. 2B). These results indicate that reporter construct microinjection did not significantly affect embryonic viability in either species.

**Figure 2.**
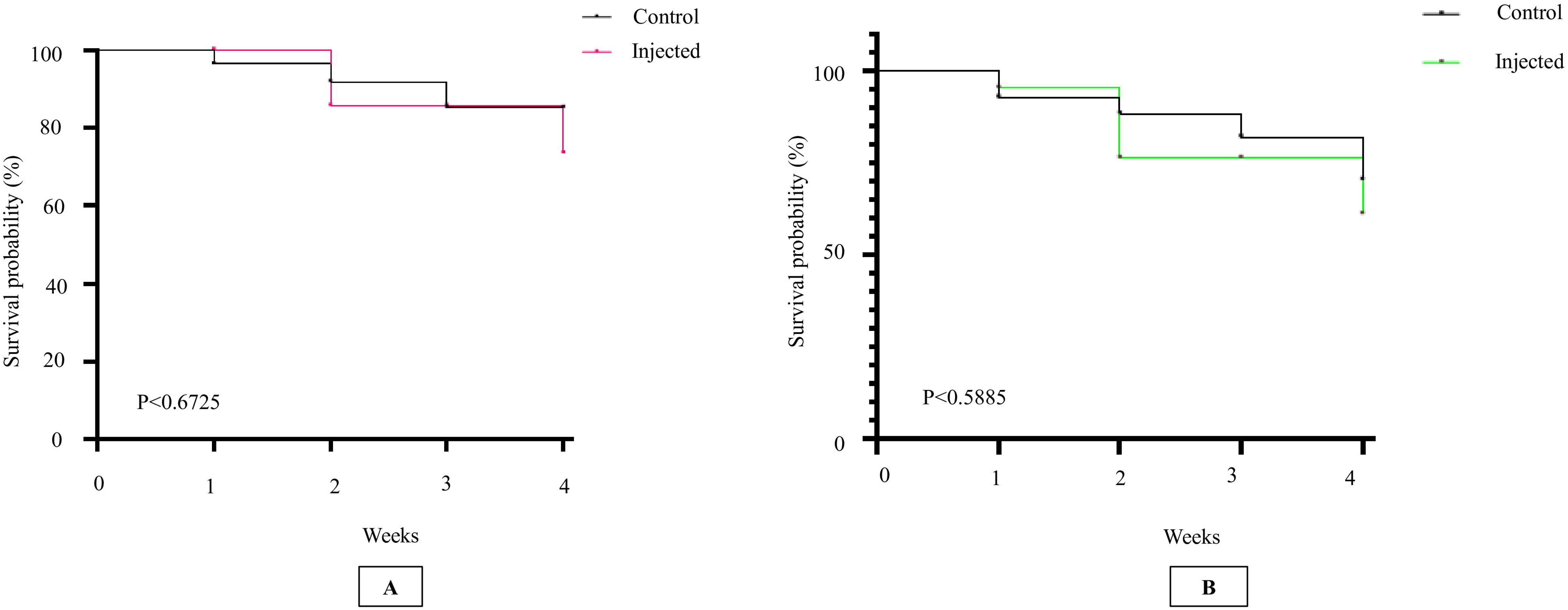
Survival analysis of embryos following reporter construct microinjection in zebrafish and sea lamprey. (A) Kaplan-Meier survival curve of zebrafish embryos comparing uninjected controls and embryos injected with the Tg(piwil1:EGFP–UTRnanos1) reporter construct (*n* = 100 embryos per group). For each group, 100 fertilized embryos were included at the beginning of the experiment and monitored longitudinally throughout development; embryos were not selected after reaching a later developmental stage. Survival probability is plotted over developmental time relative to the initial cohort. No significant difference in survival was observed between groups (log-rank test, *P* = 0.6725). (B) Kaplan–Meier survival curve of sea lamprey embryos comparing uninjected controls and embryos injected with the reporter construct (*n* = 100 embryos per group). The initial cohort consisted of 100 fertilized embryos per group, which were monitored longitudinally throughout development using the same survival assessment criteria. Survival probability is shown over developmental time relative to the initial cohort. No significant difference in survival was detected between groups (log-rank test, *P* = 0.5885).

### Germline Reporter Expression and PGC Dynamics in Zebrafish and Sea Lamprey

The Tg(piwil1–UTRnanos1) reporter exhibited dynamic germline-associated expression across developmental stages in zebrafish (Fig. 3). GFP-positive cells were first detected at the earliest stage examined, consistent with primordial germ cell (PGC) specification (Fig. 3A). During the somite stage and at 24 hpf, GFP-labeled PGCs exhibited directed migration along the embryonic body axis (Fig. 3B, C). By 48 hpf, GFP-positive cells accumulated in the presumptive gonadal region, indicating gonadal colonization (Fig. 3D). During larval development, GFP-positive germ cells persisted and expanded, with PGCs clearly localized to the developing gonadal region at 3 and 5 dpf (Fig. 3E, F). GFP-positive germ cells remained associated with the developing gonads at 10 and 15 dpf and were confined to gonadal tissue by 24 dpf (Fig. 3G-I). At 90 dpf, GFP-positive germ cells were detected within the mature gonad, including oogonia and developing oocytes, demonstrating long-term maintenance of reporter expression in differentiated germline cells (Fig. 3J, K).

**Figure 3.**
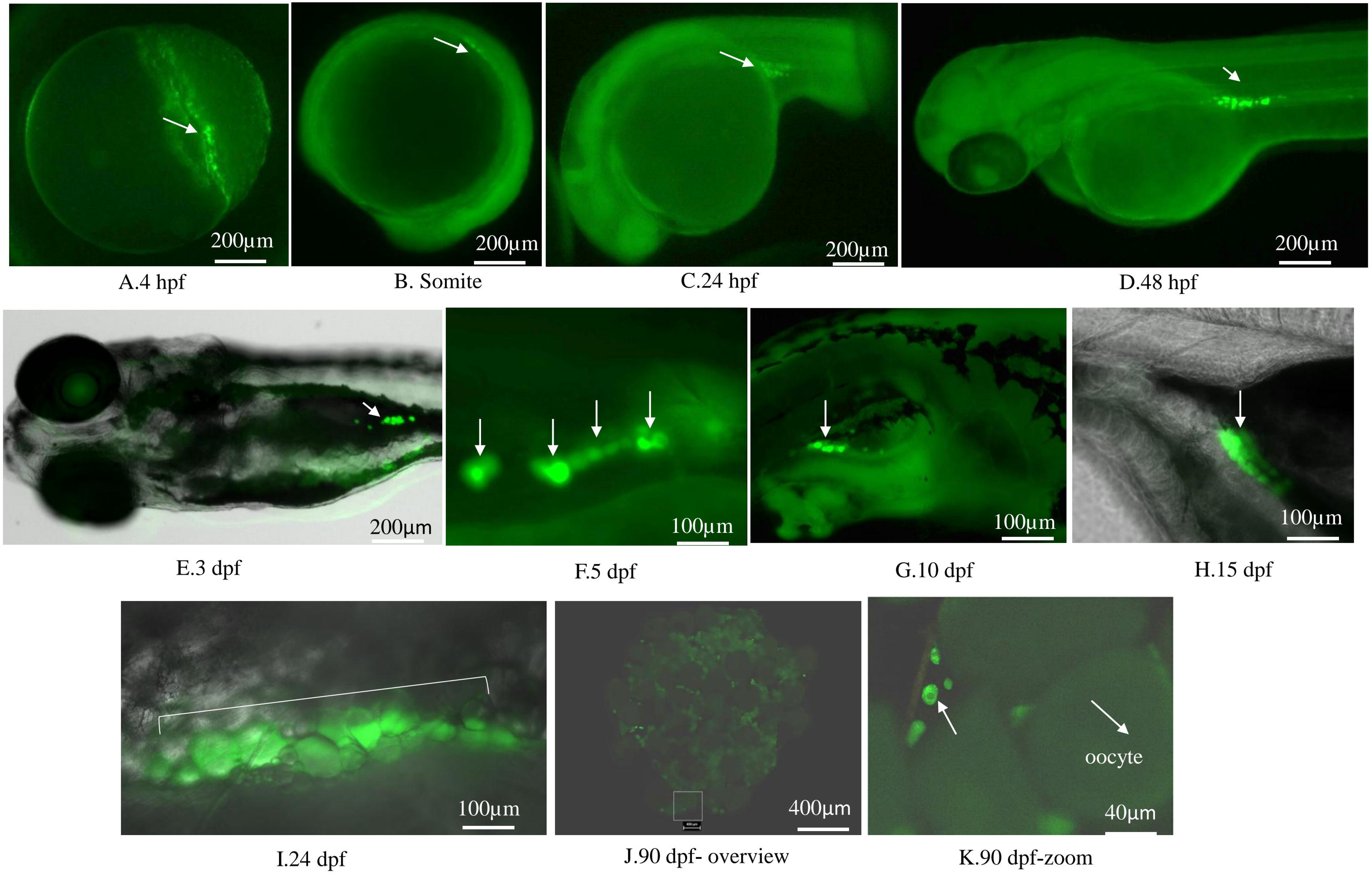
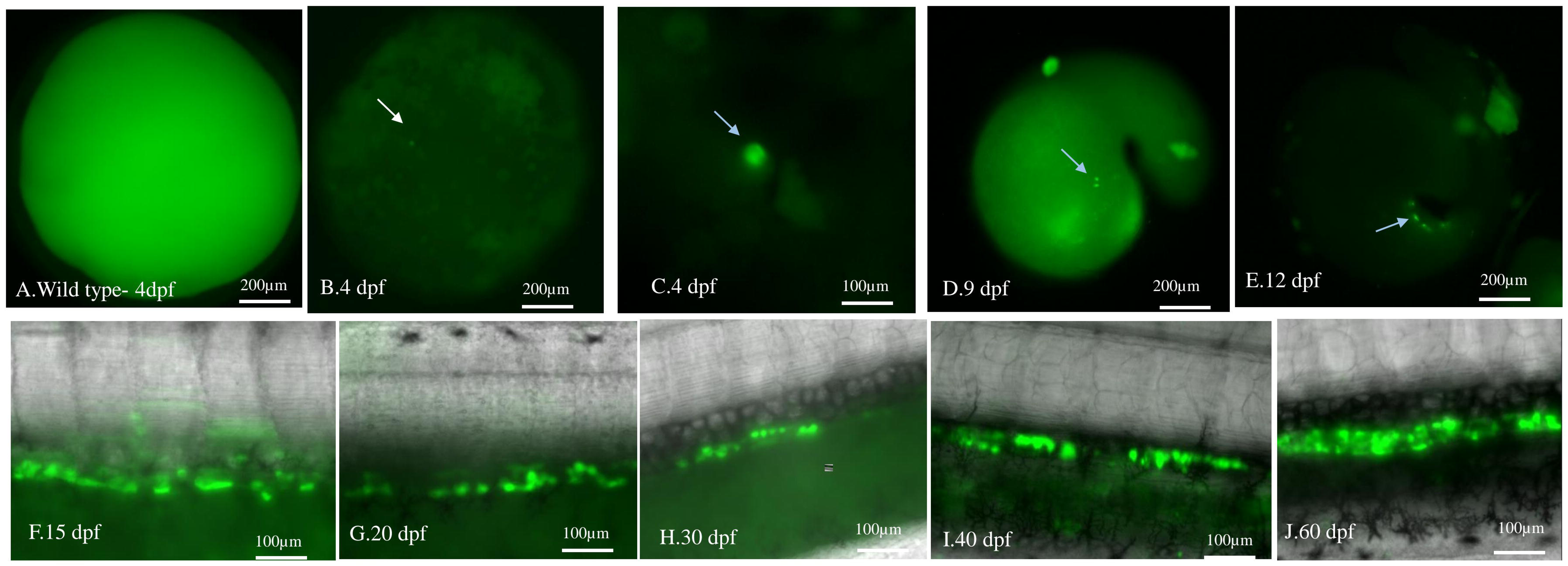
Germline reporter expression during zebrafish primordial germ cell development. (A) GFP-positive primordial germ cells (PGCs) detected at 4 hours post-fertilization (hpf). (B) PGCs during the somite stage show early migration. (C) Migrating GFP-labeled PGCs at 24 hpf. (D) Distribution of PGCs at 48 hpf during continued migration toward the gonadal region. (E-I) Progressive localization of GFP-positive germ cells during larval development at 3, 5, 10, 15, and 24 days post-fertilization (dpf), respectively, showing accumulation and persistence within the developing gonads. (J, K) Confocal imaging at 90 dpf showing GFP-positive germ cells within juvenile gonadal tissue, including oocyte structures. Merged images combine GFP fluorescence with bright-field or nuclear staining to indicate anatomical context. Scale bars are indicated in each panel.

The reporter also exhibited germline-associated expression during sea lamprey development (Fig. 4). No GFP signal was observed in wild-type controls at 4 dpf, whereas GFP-positive cells were detected in reporter-injected larvae as early as 4 dpf (Fig. 4A-C). At 9-15 dpf, GFP-positive cells increased in number and were distributed along the developing body axis, including regions adjacent to the gut (Fig. 4D-F). By 20-30 dpf, GFP-positive cells became more spatially restricted and accumulated within the presumptive gonadal region (Fig. 4G, H). At 40-60 dpf, GFP-positive germ cells remained localized within the developing gonads, indicating persistence of reporter-positive germ cells during larval development (Fig. 4I, J).

**Figure 4.**
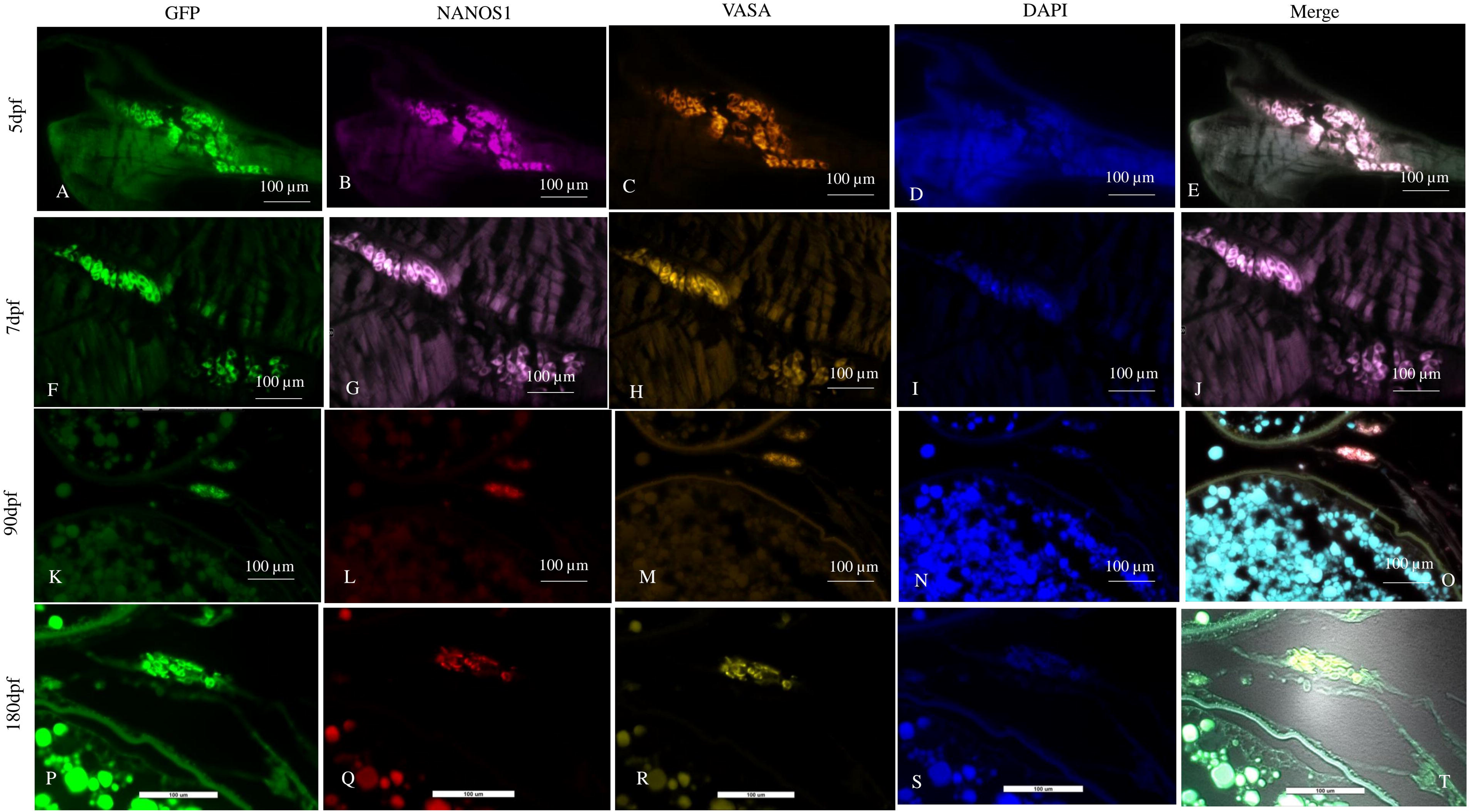
Germline reporter expression during sea lamprey larval development. (A) Wild-type larva at 4 dpf showing absence of GFP signal. (B, C) Early detection of GFP-positive primordial germ cells (PGCs) at 4 dpf. (D–F) Progressive increase and distribution of GFP-labeled germ cells at 9, 12, and 15 dpf, respectively, with cells aligning along the developing body axis and near the gut region. (G-J) Accumulation and persistence of GFP-positive germ cells at 20, 30, 40, and 60 dpf, showing localization within the presumptive gonadal region during larval development. Fluorescence images depict EGFP signal marking germline cells, with corresponding bright-field views providing anatomical context. Scale bars are indicated in each panel.

### Molecular Validation of Germline Reporter-Positive Cells in Zebrafish and Sea Lamprey

GFP-positive cells co-localized with established germline markers in zebrafish across larval and gonadal developmental stages, supporting their germline identity (Fig. 5). At 4 dpf, GFP-positive cells were detected in the developing embryo and showed spatial overlap with NANOS1 and VASA signals, consistent with early germline reporter activity (Supplementary Fig. S1A-E). At 10 dpf, GFP-positive cells remained associated with the developing gonadal region and showed continued spatial correspondence with NANOS1 and VASA expression (Supplementary Fig. S1F-J). At 5 dpf, GFP-positive cells co-localized with NANOS1 and VASA (Fig. 5A-E), and this pattern persisted at 7 dpf (Fig. 5F-J). At 90 dpf, GFP-positive germ cells were detected within the ovary, including oogonia and developing oocytes, and were organized within ovarian tissue surrounded by somatic follicular (granulosa) cell layers (Fig. 5K-O). At 180 dpf, GFP-positive germ cells remained detectable within developing gonadal tissue, demonstrating persistence of reporter expression at a later developmental stage (Fig. 5P-T). GFP-positive germ cells were also detected in the 90 dpf testis, where they were associated with seminiferous tubules and co-localized with NANOS1 (Fig. 5U-Z). The corresponding bright-field image showed the organization of the seminiferous tubules and the anatomical context of the reporter-positive germ cells (Fig. 5Z1). Together, these findings demonstrate persistent germline-associated reporter expression across larval and gonadal developmental stages in both ovarian and testicular tissues.

**Figure 5.**
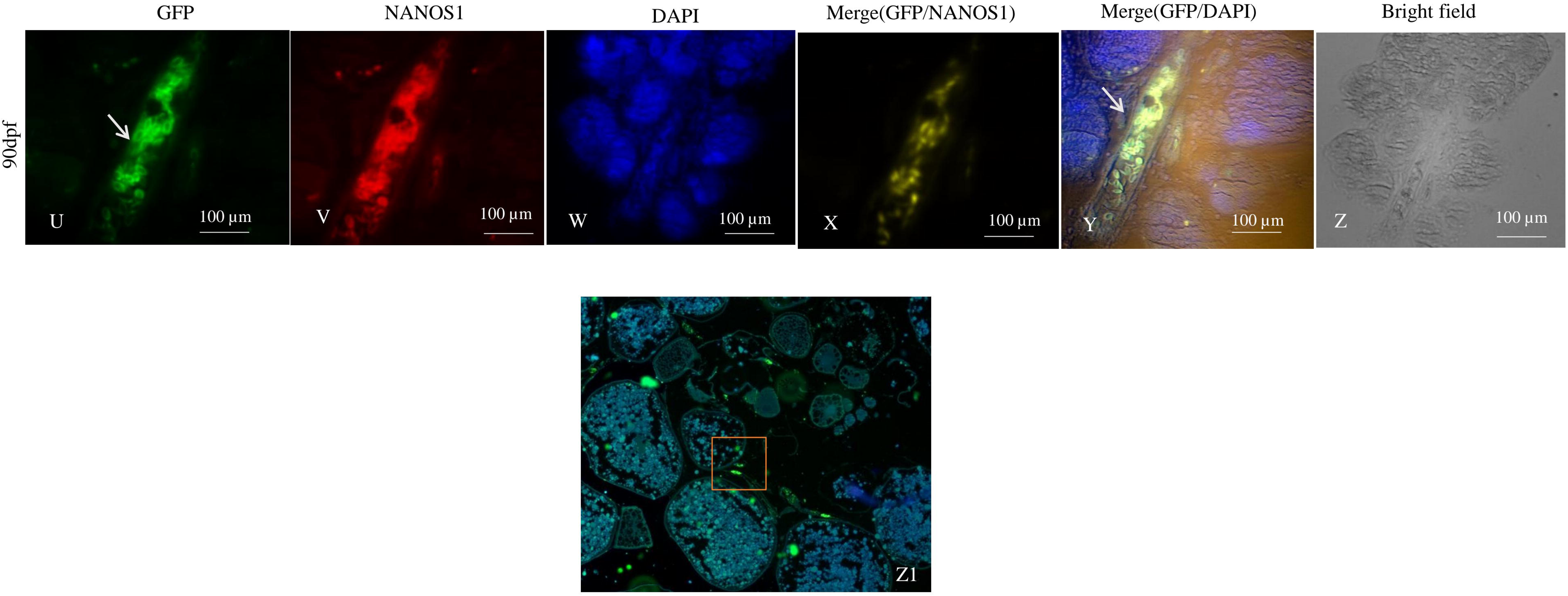
Germline-specific reporter expression and germ cell dynamics across developmental stages in zebrafish. (A-E) Representative fluorescence images at 5 days post-fertilization (5 dpf) showing expression of the Tg(piwil1:EGFP–UTRnanos1) reporter and co-localization with germ cell markers. GFP (green) marks reporter-positive germ cells, NANOS1 (red) and VASA (magenta) label germline cells, and DAPI (blue) labels nuclei. (F-J) Representative fluorescence images at 7 days post-fertilization (7 dpf) showing persistence of germline-specific reporter expression and co-localization with germ cell markers during early germ cell development. (K-O) Representative fluorescence images at 3 months post-fertilization (90 dpf) showing GFP-positive germ cells within the ovary, including oogonia and developing oocytes. Germ cells are organized within ovarian tissue and surrounded by somatic follicular (granulosa) cell layers. (P-T) Representative fluorescence images at 6 months post-fertilization (180 dpf) showing GFP-positive germ cells within the developing gonadal tissue. GFP (green) marks reporter-positive germ cells, NANOS1 (red) and VASA (magenta) label germline cells, and DAPI (blue) labels nuclei. (U-Z) Representative fluorescence and bright-field images of the testis at 3 months post-fertilization (90 dpf). (U) GFP fluorescence showing reporter-positive germ cells within the seminiferous tubules. (V) NANOS1 immunofluorescence. (W) DAPI staining showing nuclei. (X) Merged GFP and NANOS1 signals demonstrating co-localization in germ cells. (Y) Merged GFP, DAPI, and bright-field image illustrating the organization of the seminiferous tubules. (Z) Bright-field (DIC) image of the corresponding testis section. Arrows indicate GFP-positive germ cells. (Z1) Schematic model illustrating primordial germ cell (PGC) progression, germline maintenance, and differentiation into mature germ cells, highlighting germline continuity across developmental stages and transmission to the next generation. Scale bars are indicated in each panel.

In sea lamprey, GFP-positive cells formed spatially restricted clusters at Tahara stage 19 (∼20 dpf) and co-localized with the conserved germline markers NANOS1 and VASA (Fig. 6). Whole-mount imaging revealed discrete GFP-positive cell clusters within spatially restricted regions of the developing embryo (Fig. 6A, B, G, L, Q). Co-immunofluorescence showed co-localization of GFP with NANOS1 (Fig. 6C, H, R) and VASA (Fig. 6M, S), while DAPI delineated nuclear architecture (Fig. 6E, J, O, T). Merged images revealed co-localization of GFP with NANOS1 and VASA within the same cellular populations, supporting the germline identity of reporter-positive cells (Fig. 6F, K, P, U-V). GFP-positive cells remained detectable at 60 and 180 dpf and showed spatial correspondence with NANOS1- and VASA-positive cells at both stages (Supplementary Fig. S2).

**Figure 6.**
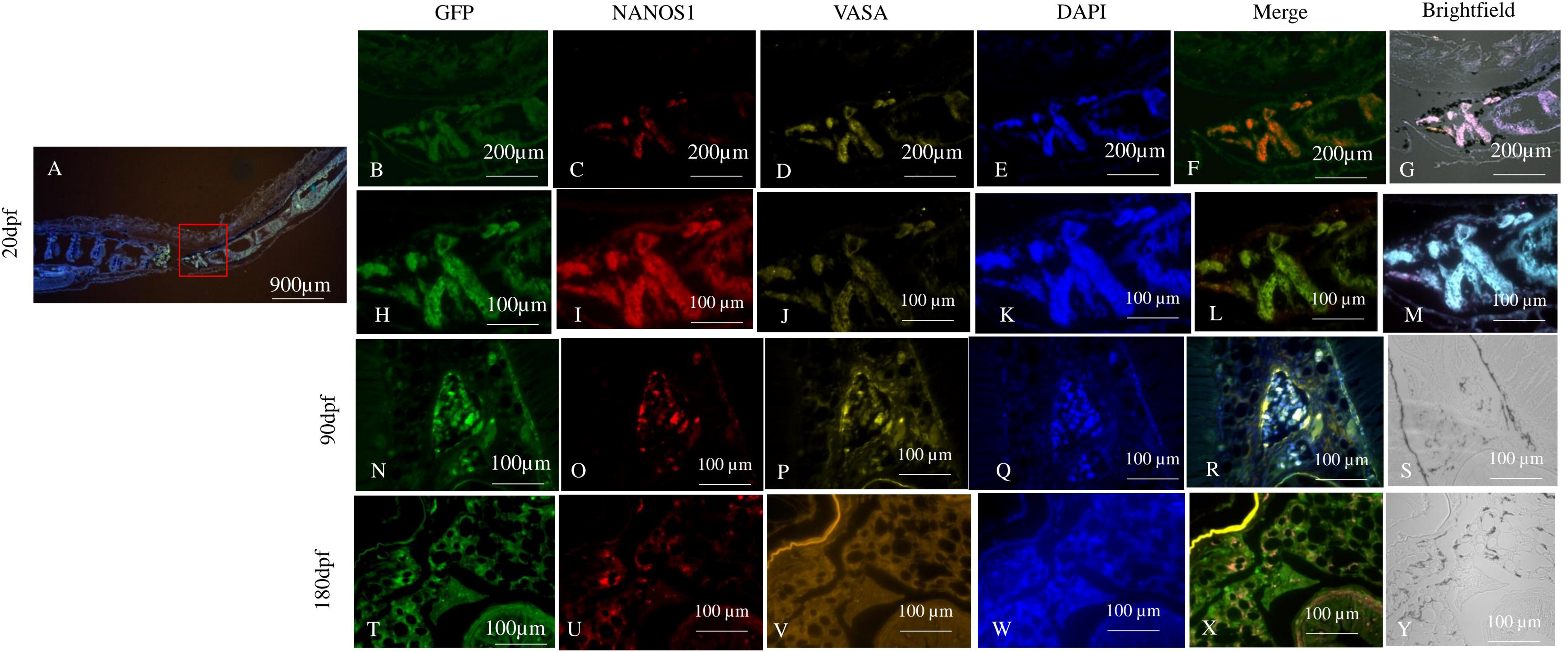
Temporal and spatial analysis of piwil1: GFP-labelled germ cells in sea lamprey. (A) Whole-mount fluorescence image of a sea lamprey larva at ∼20 dpf (Tahara stage 19). GFP-positive cells are localized to germ cell–enriched regions. The boxed area indicates the region selected for higher-magnification analysis. Scale bar, 900 µm. 20 dpf (Tahara stage 19) (B-G) Low-magnification views showing the distribution of GFP-positive cells. (H-M) High-magnification images of the same region showing cellular detail and marker co-localization. 90 dpf (N-S) GFP-positive cells and corresponding channel/merged images demonstrating maintenance of germline marker expression. 180 dpf (T-Y) GFP-positive cells and merged images showing persistence of germline identity at later stages. Channel definition (applies to H-Y): GFP (green), NANOS1 (red), VASA (magenta), and DAPI (blue); merged images show co-localization of indicated markers. GFP-positive cells consistently co-express NANOS1 and VASA across developmental stages, confirming germline identity and demonstrating conservation of germ cell markers in sea lamprey. Scale bars, 100 µm (B-Y); 200 µm where indicated.

### PGC Quantification and Sexual Differentiation

To examine the relationship between germ cell abundance and sexual differentiation in zebrafish, we analyzed PGC behavior across key developmental stages (Fig. 7). GFP-positive PGCs exhibited directed migration at 24 hpf (migration phase) and accumulated within the gonadal ridge by 48 hpf (gonadal colonization). At 5 dpf (PGC quantification stage), variation in PGC abundance among embryos began to follow a bimodal distribution, allowing classification of individuals into PGC-low and PGC-high groups (Fig. 7 A-D). Germ cells persisted in the developing gonadal region at 10-15 dpf (the gonadal differentiation stage). Fish derived from the PGC-low and PGC-high groups showed different sex-ratio distributions at 60-90 dpf, with a male-biased outcome in the PGC-low group and a female-biased outcome in the PGC-high group (Fig. 7E). Together, these observations indicate that early variation in germ cell abundance is maintained during development and is associated with divergent sexual outcomes.

**Figure 7.**
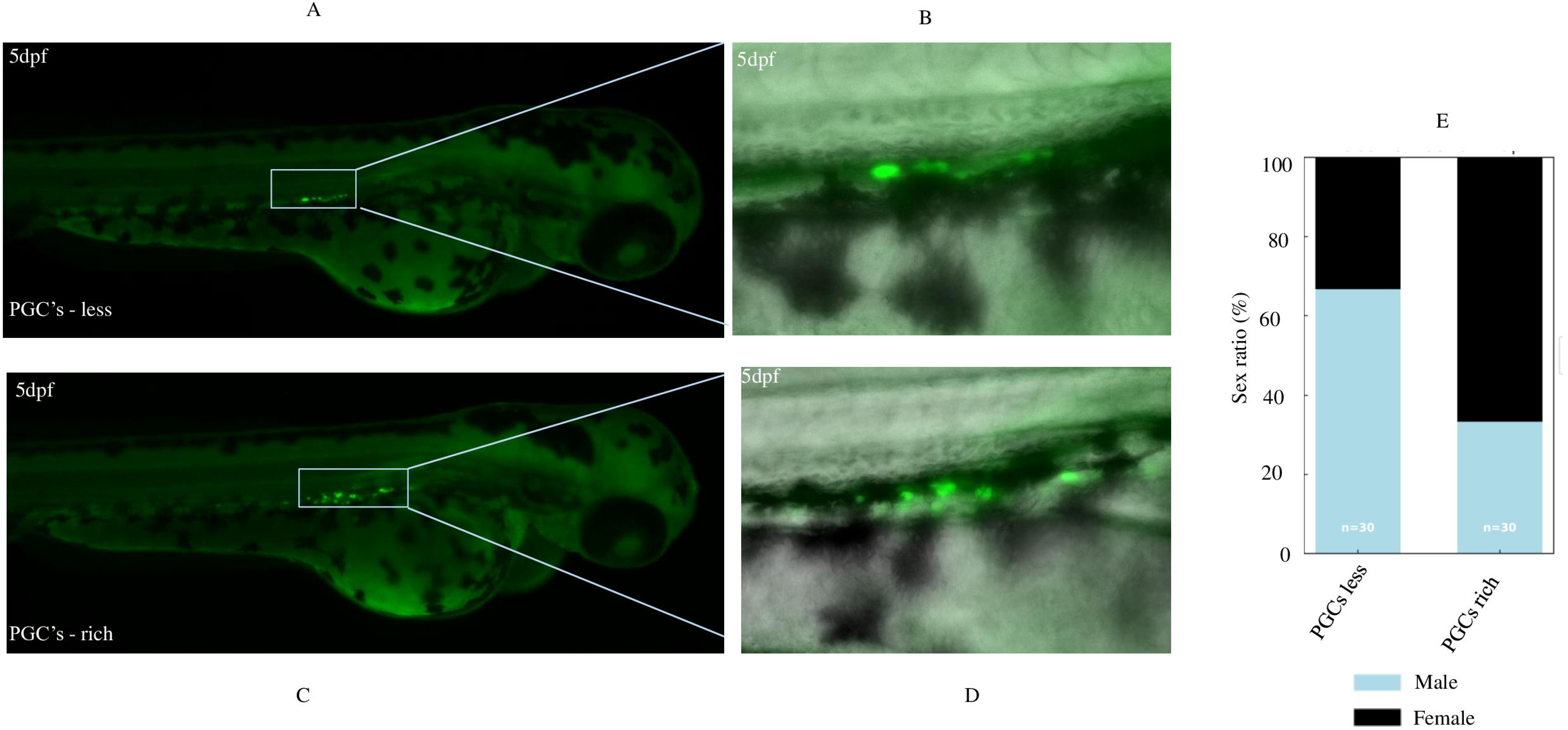
Primordial germ cell (PGC) abundance correlates with sexual development in zebrafish. (A, B) Representative fluorescence images of 5 dpf larvae with reduced PGC abundance (PGC-low group). (A) Whole larva view; (B) higher-magnification view of the gonadal region. (C, D) Representative fluorescence images of 5 dpf larvae with high PGC abundance (PGC-high group). (C) Whole larva view; (D) magnified view of the gonadal region. (E) Sex ratio at sexual maturity in fish derived from PGC-low and PGC-high groups, showing a male-biased outcome in the PGC-low group and a female-biased outcome in the PGC-high group. Arrows indicate EGFP-positive PGCs. Scale bars: 200 µm (A, C); 50 µm (B, D).

## Discussion

Understanding the mechanisms that regulate PGC specification and germline maintenance is fundamental to reproductive biology and vertebrate evolution. In this study, we developed and validated a germline-specific reporter using regulatory elements from the sea lamprey piwil1 gene coupled with the *nanos1* 3′ untranslated region, enabling visualization of germline cells in both zebrafish and sea lamprey. This approach provides a functional framework for studying germline lineages in jawless vertebrates and offers new insight into the evolutionary conservation of germline regulatory pathways.

Our findings show that regulatory elements of the sea lamprey *piwil1* gene can be used to establish a germline-specific reporter capable of marking PGCs and providing insight into conserved features of germline development across jawless and jawed vertebrates. The Tg(piwil1:EGFP-UTRnanos1) reporter produced germline-associated GFP expression in both zebrafish and sea lamprey, and reporter-positive cells co-localized with the established germline markers NANOS1 and VASA. In zebrafish, the spatial and developmental distribution of GFP-positive cells closely resembled that reported for native zebrafish *piwil1*- and *vasa*-based germline reporters, including characteristic PGC migration and gonadal localization. In sea lamprey, GFP-positive cells exhibited a corresponding progression toward the presumptive gonadal region. Together, these complementary lines of evidence support the utility of Tg(piwil1:EGFP-UTRnanos1) as a germline reporter and establish a framework for investigating germline development in a jawless vertebrate.

Selection of the sea lamprey *piwil1* regulatory region and the *nanos1* 3′ UTR was based on their complementary roles in germline regulation. The PIWI-piRNA pathway is broadly conserved and contributes to germline genome protection and maintenance (28)(Ku HY, Lin H. 2014) (29)(Zawalska M, Tarnowski M.,2025). (30)(Juliano C et al.,2011). (31, 32)(Thomson & Lin, 2009;Iwasaki et al., 2015). (34–38), whereas the *nanos* 3′ UTR provides post-transcriptional regulation that can enhance germline-restricted expression (34) (Suzuki H et al.,2010). (35, 36) (Mishima Y et al.,2006;Jin YN et al.,2018). (37)(Lee CY, et al.,2017). (13)(Hansen & Pelegri, 2021). Combining these regulatory elements therefore provided a rational strategy for developing a GSR and testing whether germline regulatory features of a cyclostome remain functional in a gnathostome model. Consistent with these findings, our comparative sequence analysis revealed strong evolutionary conservation of Piwil1 among vertebrates. The conservation of PIWI family proteins across phylogenetically distant taxa suggests that core germline regulatory mechanisms originated early in animal evolution. Previous studies have shown that the PIWI-piRNA pathway is deeply conserved and functions as a genome defense system in germ cells by preventing transposon mobilization and maintaining genomic integrity during gametogenesis (32)(Iwasaki YW et al.,2015). In zebrafish, the PGC expression pattern observed with Tg(piwil1:EGFP-UTRnanos1) closely resembles that reported for native zebrafish *piwil1*- and *vasa*-based germline reporters, with GFP-positive cells showing the characteristic distribution and developmental persistence of PGCs (48, 55)Ye D et al., 2019; Krovel AV et al., 2002).

PIWI family proteins, including Piwil1, play a central role in germline development across metazoans. Studies have demonstrated that PIWI proteins and piRNAs form specialized RNA-protein complexes that silence transposons and regulate germline stem cell maintenance and differentiation, thereby protecting the genomic stability of germline lineages (31, 32)(Thomson T and Lin H 2009; Iwasaki YW et al.,2015). In addition to their role in genome protection, PIWI proteins are widely recognized as key regulators of germline stem cell self-renewal and gametogenesis. Mutations in PIWI genes in model organisms such as Drosophila, zebrafish, and mice result in defects in germline maintenance and fertility, highlighting the conserved importance of this pathway across animals. These functions are mediated through epigenetic regulation, RNA silencing, and chromatin modification mechanisms that control gene expression during germ cell development (30, 56) (Juliano C et al.,2011; Bamezai S et al.,2012).

The development of germline reporters based on conserved regulatory elements also provides valuable tools for comparative developmental studies across vertebrates. In zebrafish, GFP-positive PGCs showed directed migration during early embryogenesis and accumulated in the gonadal region by 48 hpf, whereas in sea lamprey, GFP-positive germ cells appeared later and remained distributed along the body axis and near the gut before becoming concentrated in the presumptive gonadal region at 20-30 dpf. Thus, despite the markedly different developmental timescales, both species exhibited a broadly similar progression from early germ cell distribution and migration to gonadal localization. These findings support conservation of fundamental germline migratory and localization patterns across jawless and jawed vertebrates, while suggesting that their timing and anatomical trajectories have diverged during evolution.

The cross-species activity of the lamprey *piwil1* regulatory elements observed in this study is consistent with previous reports demonstrating that germline regulatory elements can retain functional activity across distantly related vertebrates. To test the functional capacity of lamprey regulatory elements, we validated the piwil1-based germline reporter construct in zebrafish embryos. The reporter exhibited robust and germline-specific expression during multiple developmental stages, including early germ cell specification, migration, and colonization of the gonadal ridge. These expression patterns closely mirror previously described migration routes of zebrafish primordial germ cells, which originate during early embryogenesis and migrate along defined pathways to the developing gonads. Such conserved migration dynamics highlight the value of zebrafish as a validation model for germline regulatory constructs (48)(Ye D et al.,2019).

In previous studies, the *vasa* 3′ UTR from black rockfish (*Sebastes schlegelii*) was shown to label PGCs when tested in zebrafish (*Danio rerio*) and marine medaka (*Oryzias melastigma*), supporting functional conservation of germline regulatory mechanisms across teleosts (57, 58) (McEwen GK et al.,2009; Zhou L et al.,2020). Our data further support the utility of the present reporter for studying conserved germline lineage dynamics across distant vertebrate groups.

We found that embryos with reduced PGC numbers showed a male-biased sex ratio, whereas individuals with higher germ cell abundance tended to develop as females. Similar relationships between PGC abundance and gonadal differentiation have been reported in other teleost species, where germ cell migration dynamics and early PGC number are closely associated with sex development (38)(Chu WK et al., 2025). Interestingly, early work in lamprey suggested that PGC number may influence reproductive potential, with variations in germ cell abundance correlated with fecundity in cyclostomes (21, 22, 59) (Yasmin T et al.,2022; Tezak B et al., 2023; Hardisty & Cosh, 1966). Our study therefore provides a framework for investigating whether the relationship between PGC abundance and sexual differentiation is also present in sea lamprey. Because the Tg(piwil1-UTRnanos1) reporter enables visualization and quantification of germline cells, it provides an experimental approach to test whether the association between PGC abundance and sex differentiation observed in zebrafish is conserved in this jawless vertebrate. These findings are consistent with previous reports demonstrating that germ cell quantity influences sex differentiation in zebrafish. Experimental depletion of PGCs has been shown to bias development toward testis formation, whereas increased PGC proliferation promotes ovarian differentiation. Thus, PGCs are not merely passive participants in gonadal development but may actively contribute to sexual fate decisions (23, 60). (Slanchev K et al.,2005; Tzung KW et al.,2015)

Our finding that reduced PGC abundance was associated with a male-biased sex ratio, whereas higher PGC abundance was associated with a female-biased outcome, further supports a relationship between PGC abundance and gonadal sex differentiation during early development. Rather than providing new evidence for this relationship in zebrafish, our findings demonstrate that the Tg(piwil1-UTRnanos1) reporter can be used to visualize and quantify PGC abundance and provide a framework for testing whether the relationship between germ cell dynamics and sexual differentiation is conserved in sea lamprey.(23)(Tzung KW et al.,2015). Our reporter provides a framework for testing whether this relationship is conserved in jawless vertebrates.

Beyond its utility for tracking germline development, the reporter provides a platform for investigating the relationship between germ cell dynamics and sexual development. (61) (Timoshevskiy VA et al.,2025). This may be particularly valuable in vertebrate species in which sex determination and differentiation involve genetic sex-determining mechanisms (GSD), environmental influences (ESD), or prolonged and developmentally labile periods of sexual differentiation. Longitudinal visualization of PGC abundance, migration, and gonadal colonization could allow these processes to be examined in relation to subsequent gonadal fate and sex differentiation. Comparative application of the reporter or related germline reporters across species with contrasting sex-determination systems may therefore help determine whether relationships between germ cell dynamics and sexual fate represent conserved features of vertebrate development or lineage-specific adaptations.

A defining feature of PGC development is extensive epigenetic reprogramming that resets the germline epigenome and establishes the developmental potential required for transmission of the germline to the next generation. During this process, genome-wide DNA methylation patterns are extensively remodeled, and parental imprinting marks are erased and subsequently established during gametogenesis. Although these processes have been extensively characterized in mammals, how epigenetic reprogramming is regulated in early-diverging vertebrates such as lampreys remains largely unknown (49–54)(Lee SM, Zhu Q et al.,2021; Surani MA., Singh A et al.,2023; Messerschmidt DM et al.,2014; Angeloni A et al.,2024). The germline-specific reporter developed here provides a new experimental framework for identifying and following germline cells in sea lamprey and may facilitate future investigation of the molecular and epigenetic mechanisms underlying germline development and maintenance in this early-diverging vertebrate lineage.

This reporter system, therefore, provides an important molecular tool for investigating germline biology in jawless vertebrates and offers new opportunities to explore the evolutionary origins of vertebrate reproductive systems. By enabling visualization and quantification of germline cells in sea lamprey, the reporter also provides a foundation for future studies of germline-targeted genetic technologies, including the development and evaluation of gene-drive strategies for genetic management of invasive sea lamprey populations.

## Data Availability Statement

The sequence of the sea lamprey *piwil1*-GFP-*nanos1* reporter construct is available in GenBank under accession number PZ890288.

## Acknowledgements

The authors acknowledge the Great Lakes Fishery Commission (GLFC) for providing facilities and equipment that supported this research. We extend our special thanks to Nicholas Johnson for research support and collaboration, Kian Buckowski for technical assistance, and Jacob Kimmel and Kristen Lounsbury for their assistance in maintaining the zebrafish and sea lamprey facilities. We sincerely thank Dr. Robb Krumlauf, Dr. Hugo Parker, and Dr. Marianne Bronner and their colleagues at the Stowers Institute for Medical Research and Caltech for developing the lamprey reporter technology and providing the Hugo’s Lamprey Construct (HLC) vector backbone used in this study.

## Authors’ contributions

**SS**: Conceptualization, Methodology, Investigation, Formal analysis, Data curation, Visualization, Writing - original draft, Writing - review and editing. **YWCD**: Methodology, Investigation, Resources, Writing - review and editing. **TJB**: Investigation, Resources, Writing - review and editing. **JS**: Methodology, Investigation, Resources, Writing. **WL**: Conceptualization, Supervision, Project administration, Funding acquisition, Writing - review and editing.

---

**Table 1.** Comparative germline tools across vertebrates.

| S.No | Model organism | Germline marker/tool | Strategy | Key limitation | Reference |
| --- | --- | --- | --- | --- | --- |
| 1 | Zebrafish | *vasa*, *nanos* reporters | Transgenesis (Tol2) | Well-established | (62, 63)Suster ML et al.,2009; Kosaka K et al.,2007 |
| 2 | Mouse | PGCLC system | *In vitro* differentiation | Not *in vivo* lineage tracking | (64) Li L et al.,2025 |
| 3 | Xenopus | Germ plasm-based markers | Preformation | Limited genetic tools | (65)Seervai RN, Wessel GM 2013 |
| 4 | Teleosts (general) | Germline fluorescent reporters | Promoter + UTR systems | Species-specific variation | (66) Knaut H et al.,2002 |
| 5 | Sea lamprey | No stable reporter (previously)<br>*piwil1*: EGFP-UTRnanos1 | Cross-species + transgenesis | First functional tool | Present study |

**Supp Table 1.**
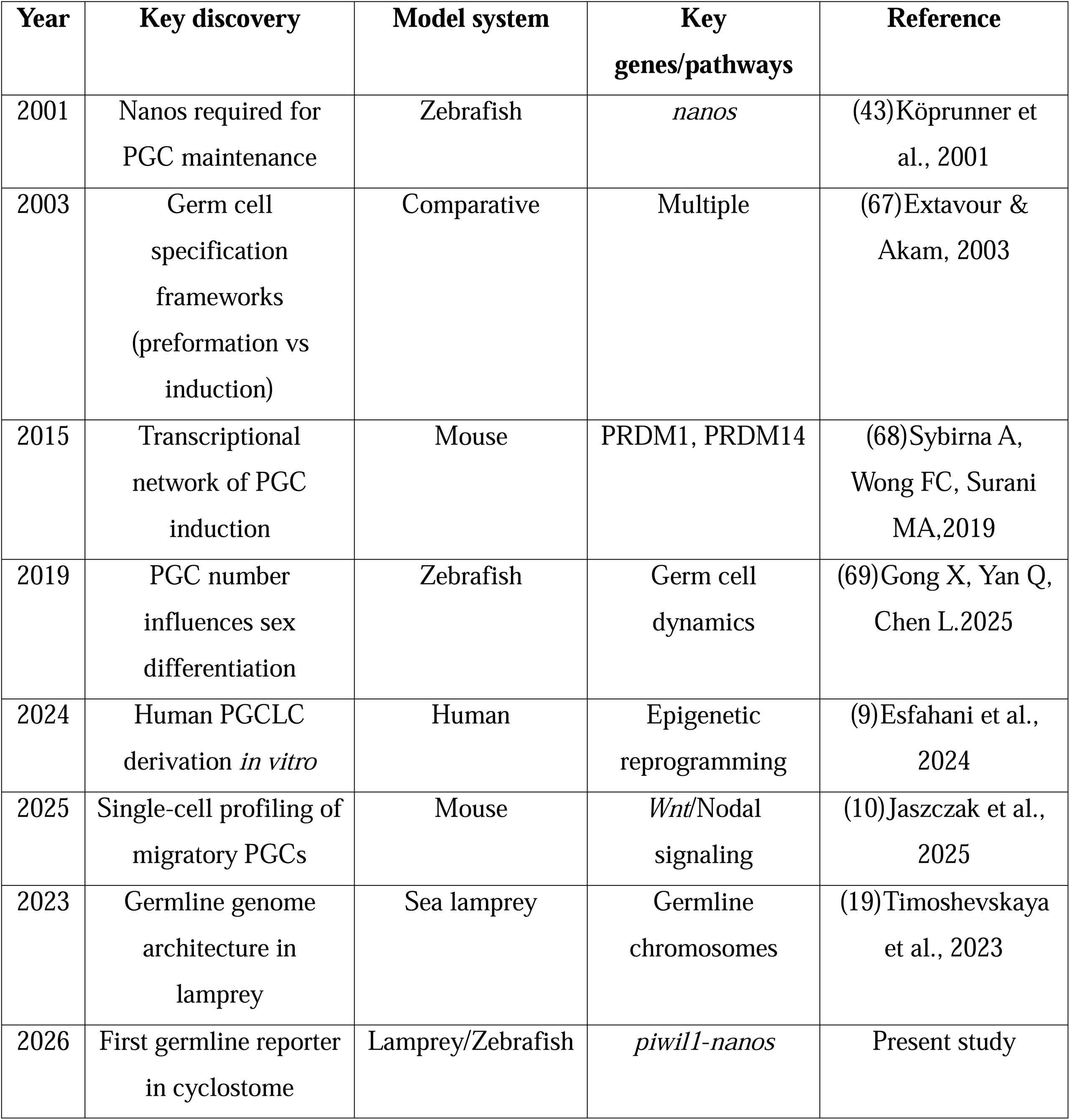
Key milestones in vertebrate PGC research.

