## Supplementary Figure S1 and S2 for "A germline-specific reporter reveals conserved primordial germ cell developmental dynamics between jawless and jawed fish species"

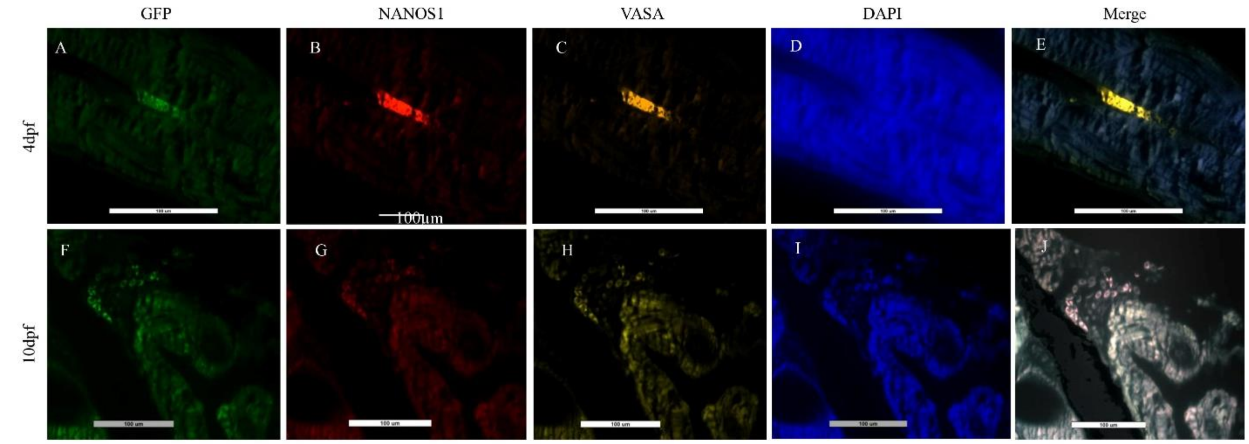


**Supplementary Figure S1. Temporal and spatial analysis of Tg(piwil1:EGFP–UTRnanos1) reporter-positive germ cells in zebrafish.**

(A-E) Representative fluorescence images of zebrafish at 4 dpf, showing GFP reporter expression and germline-marker expression. (A) GFP fluorescence identifying reporter-positive cells. (B) NANOS1 immunofluorescence. (C) VASA immunofluorescence. (D) DAPI staining showing nuclei. (E) Merged image showing the spatial relationship among GFP, NANOS1, VASA, and DAPI signals. (F-J) Representative fluorescence images of zebrafish at 10 dpf, showing persistence and spatial distribution of reporter-positive germ cells during early gonadal development. (F) GFP fluorescence. (G) NANOS1 immunofluorescence. (H) VASA immunofluorescence. (I) DAPI staining. (J) Merged image showing the spatial relationship among GFP, NANOS1, VASA, and DAPI signals. Scale bars, 100 μm.

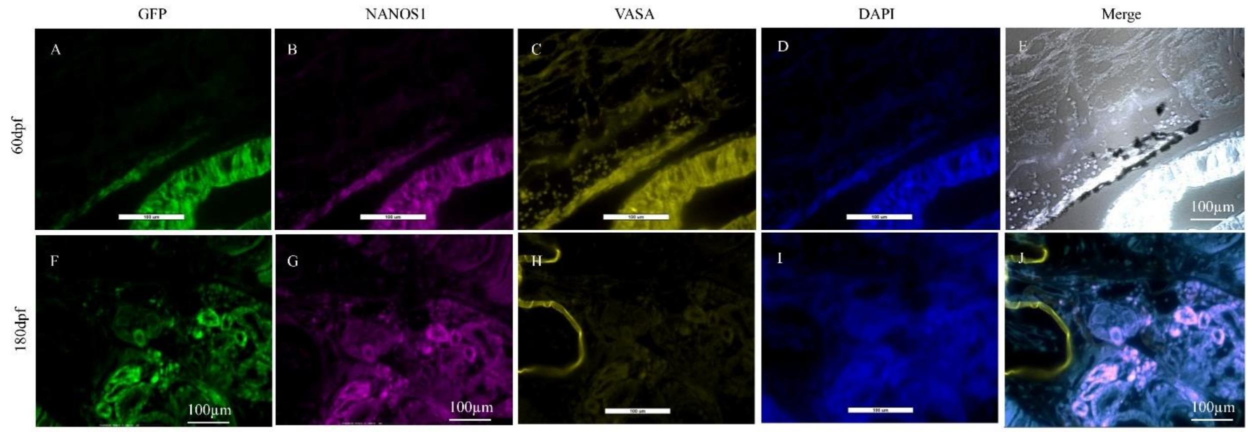


**Supplementary Figure S2. Temporal and spatial analysis of Tg(piwil1:EGFP–UTRnanos1) reporter-positive germ cells in sea lamprey.**

(A-E) Representative fluorescence images of sea lamprey at 60 dpf, showing GFP reporter expression and germline-marker expression. (A) GFP fluorescence identifying reporter-positive cells. (B) NANOS1 immunofluorescence. (C) VASA immunofluorescence. (D) DAPI staining showing nuclei. (E) Merged image showing the spatial relationship among GFP, NANOS1, VASA, and DAPI signals. (F-J) Representative fluorescence images of sea lamprey at 180 dpf, showing persistence of reporter and germline-marker expression during later gonadal development. (F) GFP fluorescence. (G) NANOS1 immunofluorescence. (H) VASA immunofluorescence. (I) DAPI staining. (J) Merged image showing the spatial relationship among GFP, NANOS1, VASA, and DAPI signals. Scale bars, 100 µm.
